# Evaluation of *staA*, *viaB* and *sopE* genes in *Salmonella* detection using conventional polymerase chain reaction (PCR)

**DOI:** 10.1101/2020.06.22.164491

**Authors:** Francis Kariuki, Pauline Getanda, Atunga Nyachieo, Gerald Juma, Peter Kinyanjui, Joseph Kamau

**Affiliations:** Department of Biochemistry, University of Nairobi, P.O. Box 30197-00100, Nairobi, Kenya; Molecular Biology Unit, Institute of Primate Research, P.O. Box 24481-00502, Nairobi, Kenya

## Abstract

Typhoid fever is caused by the bacteria *Salmonella enterica* subspecies *enterica* serovar Typhi (*S*. Typhi) and remains a significant health problem in many developing countries. The lack of adequate diagnostic capabilities in these poor resource settings have contributed greatly in making typhoid fever endemic in these regions. Reliable and inexpensive diagnostic tests are needed to improve the management of this disease burden. This study evaluated the ability of *staA, viaB* and *sopE* genes to detect *Salmonella* spp. Conventional polymerase chain reaction (PCR) amplification of *staA*, *viaB* and *sopE* genes of *Salmonella* was used to detect and differentiate between the three most prevalent *Salmonella* spp. in Kenya (*S*. Typhi, *S.* Typhimurium and *S.* Enteritidis). The *staA* primers (StaA-Forward / StaA-Reverse) and *viaB* primers (vi-Forward / vi-Reverse) were found to be specific only for the different strains of *S*. Typhi, producing PCR products of 585 bp and 540 bp respectively. No amplification was observed with *S.* Typhimurium, *S.* Enteritidis, *E*. coli and *S*. boydii bacterial strains. The *sopE* primers (SopE-Forward / SopE-Reverse) was demonstrated to be specific for all *Salmonella* spp. producing a 465 bp PCR product with no amplification observed with the *E*. coli and *S*. boydii bacterial strains. Conventional PCR using these *staA* and *viaB* primers for detection of *S*. Typhi shows great potential for diagnosis of typhoid fever however, further studies need to be carried out with actual food samples and human samples (blood, stool or saliva) to determine the effectiveness of this method in the detection of common Salmonella spp. in Kenya.

**Author summary:** Typhoid fever is a severe disease caused by the bacteria *Salmonella* Typhi (*S*. Typhi) and is a significant health problem in many developing countries. The lack of adequate diagnostic capabilities in poor resource settings common in most public health facilities in Kenya and Africa in general, hinder prompt diagnosis of typhoid fever. Currently, the available diagnostic tests are often expensive and more so not readily available in most resource poor endemic areas. This has often led to misdiagnosis of the disease, thereby delaying appropriate treatment and making typhoid fever widespread in most resource poor areas. This study examines the ability of three different genes to detect and differentiate between the three most prevalent *Salmonella* strains in Kenya using a readily available and widely used genetic test known as conventional polymerase chain reaction (PCR). This research found that *staA* and *viaB* genes were specific only for *S*. Typhi, while the *sopE* gene was specific for all *Salmonella* strains. Consequently, conventional PCR using these *staA* and *viaB* genes for detection of *S*. Typhi shows great potential to be used as a readily available diagnostic tool to detect the presence of the *S*. Typhi organism in individuals or foods sample in Kenya.

## Introduction

Microbial food borne diseases are widespread and have resulted in considerable economic losses in many parts of the world particularly the low-income developing countries of South East Asia, Africa, and Latin America [1,2]. *Salmonella* is a major foodborne pathogen responsible for a large number of food-poisoning cases in humans both in the developed and developing countries. For example, *Salmonella* accounts for approximately 65% and 11% of food poisoning cases in France [3] and the United States [4] respectively. Additionally, the costs associated with *Salmonella* infections are also remarkably high. For example, according to the World Health Organization (WHO), the *Salmonella* associated costs in the United States of America are estimated at US$ 3.6 billion annually [5].

In the sub-Saharan Africa, the disease burden is high and the cost constraints prevent the creation of robust surveillance programs and the widespread use of newer, more effective but expensive diagnostic tool, thus making *Salmonella* incidences endemic [6]. Large outbreaks of *Salmonella* infections have been associated with poor sanitation and poor hygiene conditions [7]. Transmission of *Salmonella* to humans has been linked to numerous sources, including contaminated and uncooked poultry and poultry products, meat, milk and other dairy products, pork, fresh vegetables and fruits as well as contaminated water and contact with infected animals [8,9].

Typhoid fever is a bacterial disease, caused by the typhoidal *Salmonella* serovar *S.* Typhi and is transmitted through the ingestion of food or drink contaminated by the faeces or urine of infected people or carriers. It is a significant health problem in many developing countries. Worldwide, an estimated 17 million cases [10] occur annually with most of the disease burden occurring among citizens of low-income countries, particular those in South East Asia, Africa, and Latin America. [1]. To avoid severe complications or even the loss of life because of salmonellae infections, definite and accurate diagnosis and treatment need to be initiated as soon as the onset symptoms of the infection begin to manifest. However, the lack of adequate diagnostic capabilities in poor resource settings common in most public health facilities in Kenya and Africa in general, hinder prompt diagnosis of Salmonellae infections particularly typhoid fever. This has often led to misdiagnosis of the disease, thereby delaying appropriate treatment and making typhoid fever endemic in most resource poor areas.

To improve accurate and early detection of *Salmonella* Typhi (S. Typhi) in Kenya, we tested the ability of three pairs of primers targeting three different genes: SopE invasion-associated secreted protein (Gene ID: 1250812), ViaB region DNA for Vi antigen (GI: 426443) and StaA fimbrial protein (Gene ID: 1246701) to detect and differentiate *S*. Typhi from other salmonella serovars and closely related disease causing bacteria.

## Results

### Genomic Products from DNA Extraction

An overnight (16 hrs. @ 37 °C) 5ml culture of each of the bacterial isolates was harvested and DNA extracted using the standard phenol-chloroform method. The genomic DNA products extracted from the twelve (12) bacterial isolates, was analyzed and confirmed on a 1% agarose gel (Fig 1).

**Figure 1.**
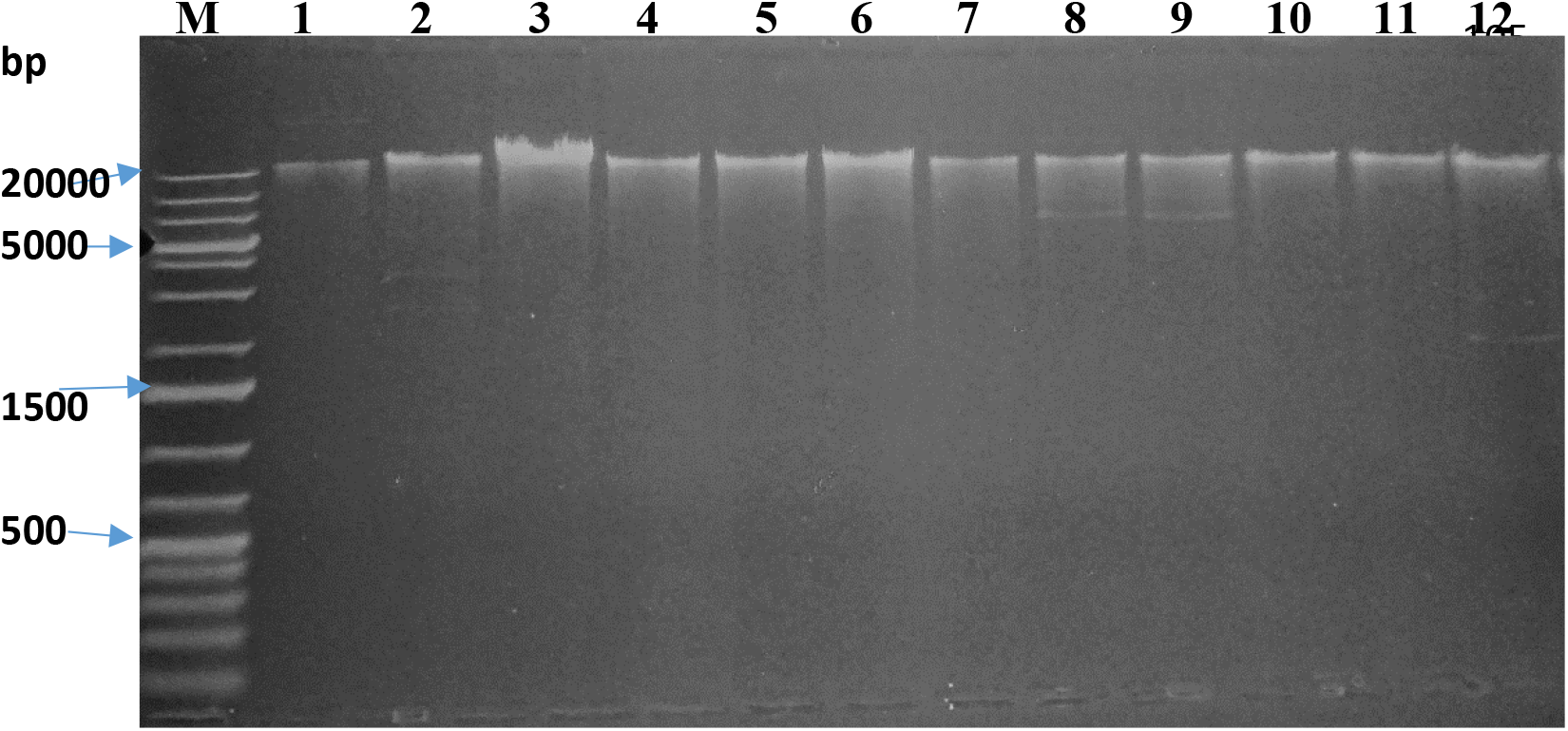
1 % Agarose gel of Genomic DNA products isolated from various bacterial isolates. Lanes 1-12 represent DNA isolated from various bacteria using the standard phenol-chloroform method. M - 1kb DNA ladder, 1-*E.* coli, 2 - *S.* boydii, 3 - *S.* Typhimurium, 4 - *S*. Typhi (s), 5 - *S*. Typhi (i), 6 - *S*. Typhi (r), 7 - *S*. Typhimurium (s), 8 - *S*. Typhimurium (i), 9 - *S*. Typhimurium (r), 10 - *S*. Enteritidis (s), 11 - *S*. Enteritidis (i), 12 - *S*. Enteritidis (r)

### Evaluation of the primers

The specificity of each set of primers (table 2) was tested against each of the bacterial isolates (table 1). The 16s rRNA primers (Minf / Minr) was able to confirm the genus *Salmonella*. Amplification occurred for all the tested *Salmonella* spp. resulting in a 402 bp pcr product. No amplification was observed from the *E*. coli, *S*. boydii bacterial strains using the same primers (figure 2).

**Table 1.** List of bacterial isolates used in the study

|  |  |  |  |  |  |  |  |
| --- | --- | --- | --- | --- | --- | --- | --- |
| 1 | *E. coli* ATCC 25922 | 4 | *S. Typhi* (S) | 7 | *S. Typhimurium* (S) | 10 | *S. Enteritidis* (S) |
| 2 | *S. boydii* ATCC 9207 | 5 | *S. Typhi* (I) | 8 | *S. Typhimurium* (I) | 11 | *S. Enteritidis* (I) |
| 3 | *S. Typhimurium* ATCC 14022 | 6 | *S. Typhi* (R) | 9 | *S. Typhimurium* (R) | 12 | *S. Enteritidis* (R) |

**Table 2.**
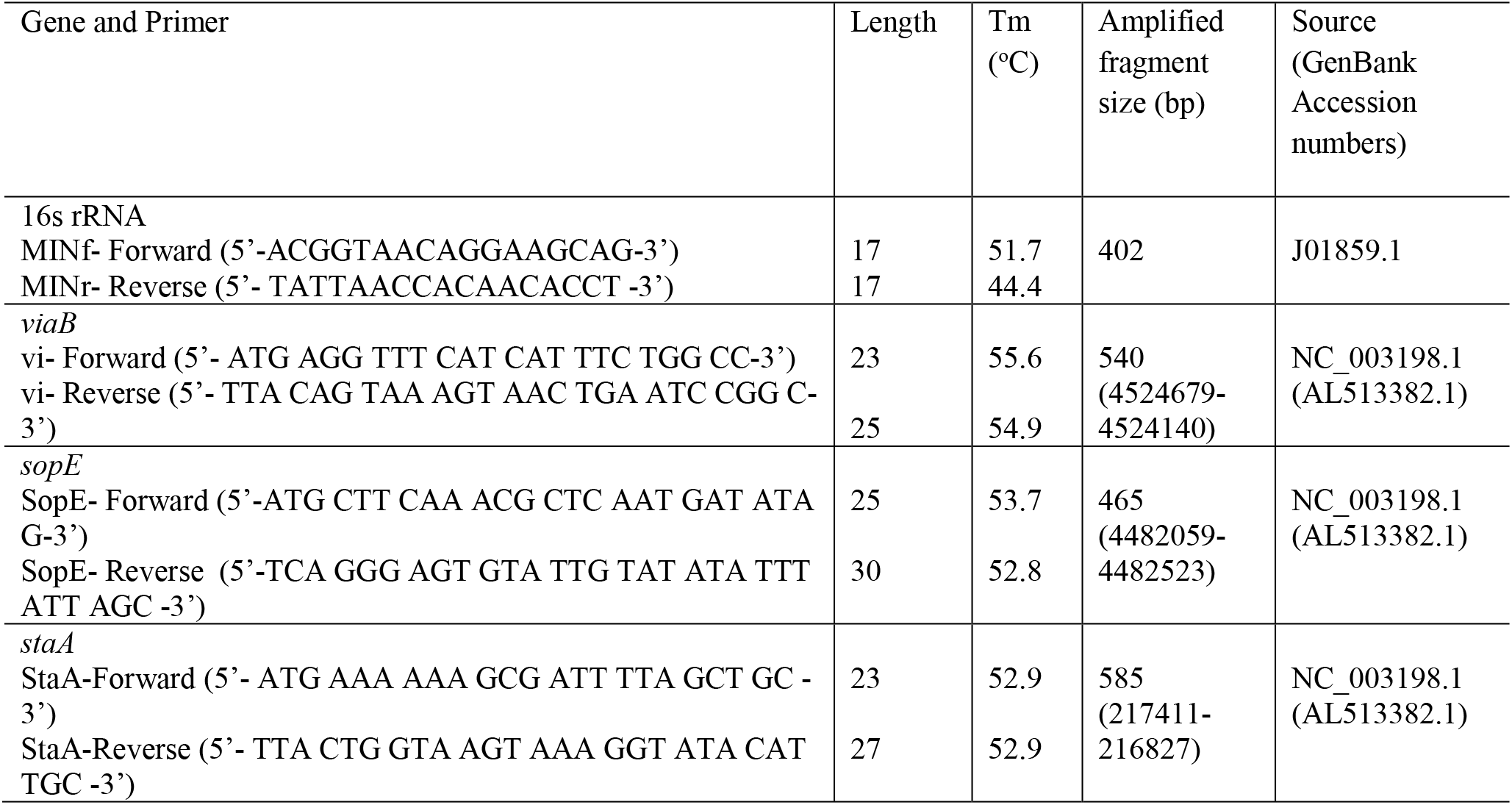
List of primer used in identification and differentiation of salmonella bacteria

**Figure 2.**
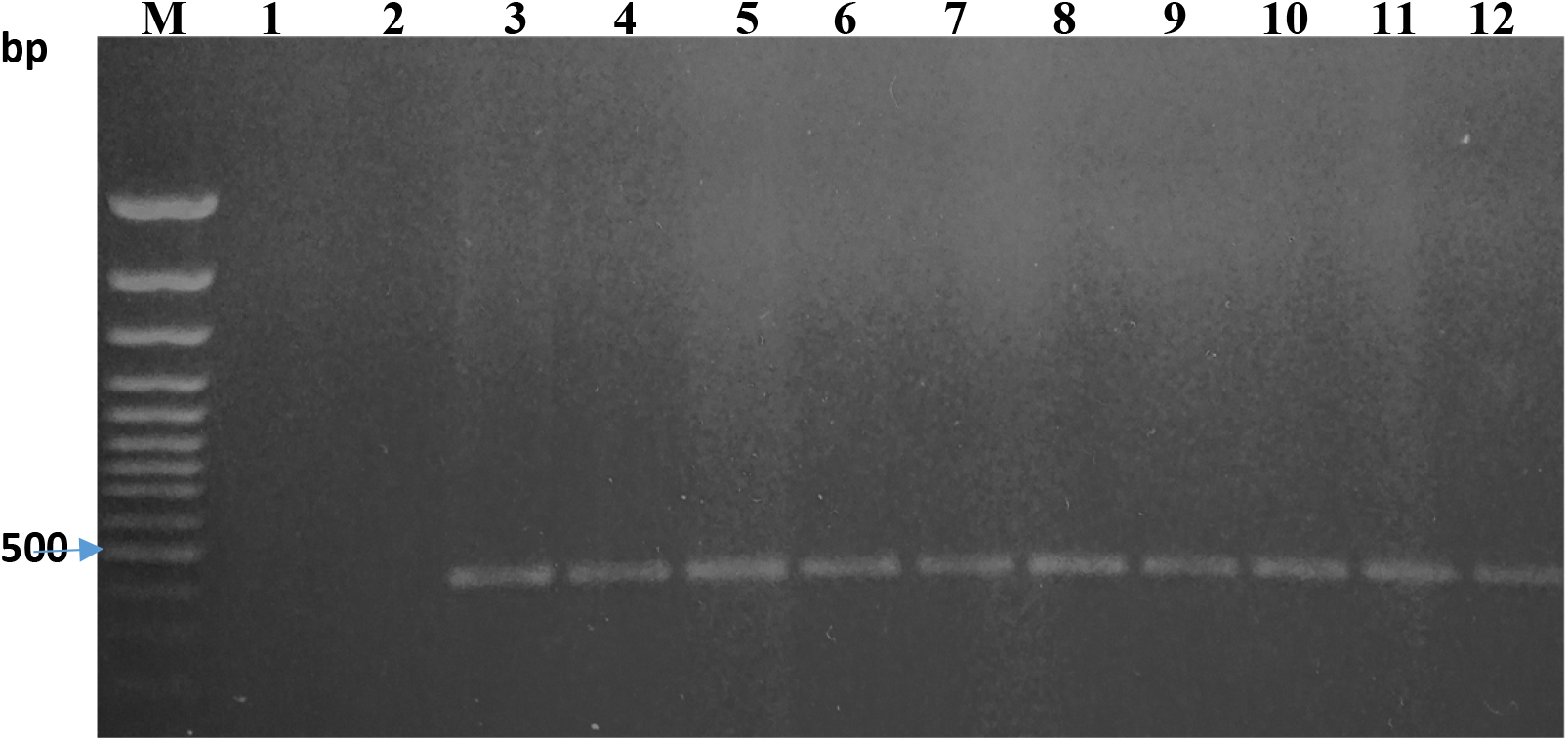
1% Agarose gel of the PCR products using 16S rRNA target primers and DNA templates extracted from various bacterial isolates. (Expected product size is 402 bp). M (100 bp DNA ladder). Lane 1-12 (PCR product of various bacteria using 16S rRNA target primers). 1-*E.* coli, 2 - *S.* boydii, 3 - *S.* Typhimurium, 4 - *S*. Typhi (s), 5 - *S*. Typhi (i), 6 - *S*. Typhi (r), 7 - *S*. Typhimurium (s), 8 - *S*. Typhimurium (i), 9 - *S*. Typhimurium (r), 10 - *S*. Enteritidis (s), 11 - *S*. Enteritidis (i), 12 - *S*. Enteritidis (r)

The *staA* primers (StaA-Forward / StaA-Reverse) and *viaB* primers (vi-Forward / vi-Reverse) were found to be specific only for the different strains of *S*. Typhi, with amplification resulting in a 585 bp (figure 3) and 540 bp (figure 4) PCR product respectively. No amplification was observed with *S.* Typhimurium, *S.* Enteritidis, *E*. coli and *S*. boydii bacterial strains.

**Figure 3.**
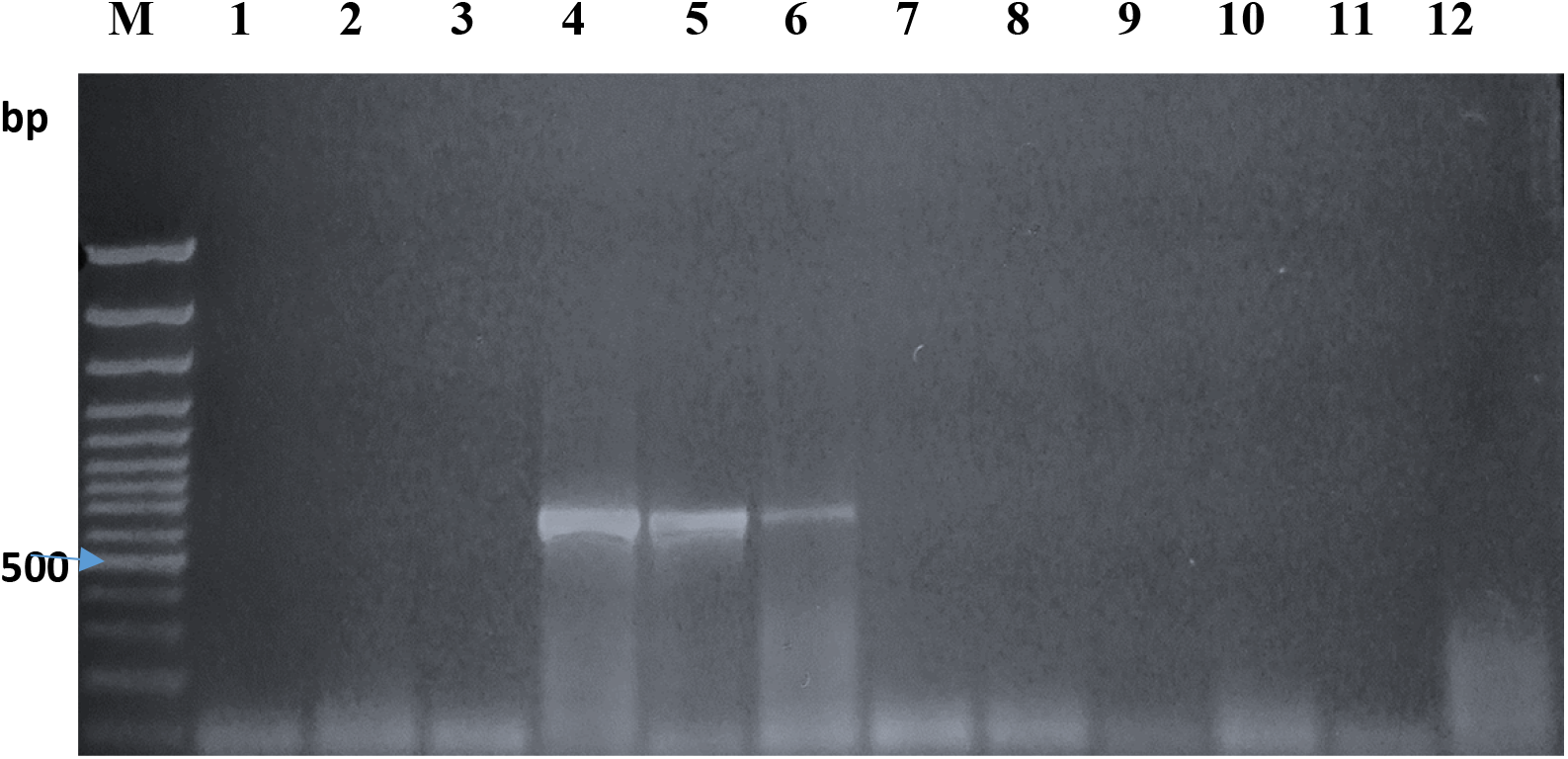
1% Agarose gel of the PCR products using *staA* target primers and DNA templates extracted from various bacterial isolates. (Expected product size is 585 bp). M (100 bp DNA ladder). Lane 1-12 (PCR product of various bacteria using *staA* target primers). 1-*E.* coli, 2 - *S.* boydii, 3 - *S.* Typhimurium, 4 - *S*. Typhi (s), 5 - *S*. Typhi (i), 6 - *S*. Typhi (r), 7 - *S*. Typhimurium (s), 8 - *S*. Typhimurium (i), 9 - *S*. Typhimurium (r), 10 - *S*. Enteritidis (s), 11 - *S*. Enteritidis (i), 12 - *S*. Enteritidis (r).

**Figure 4.**
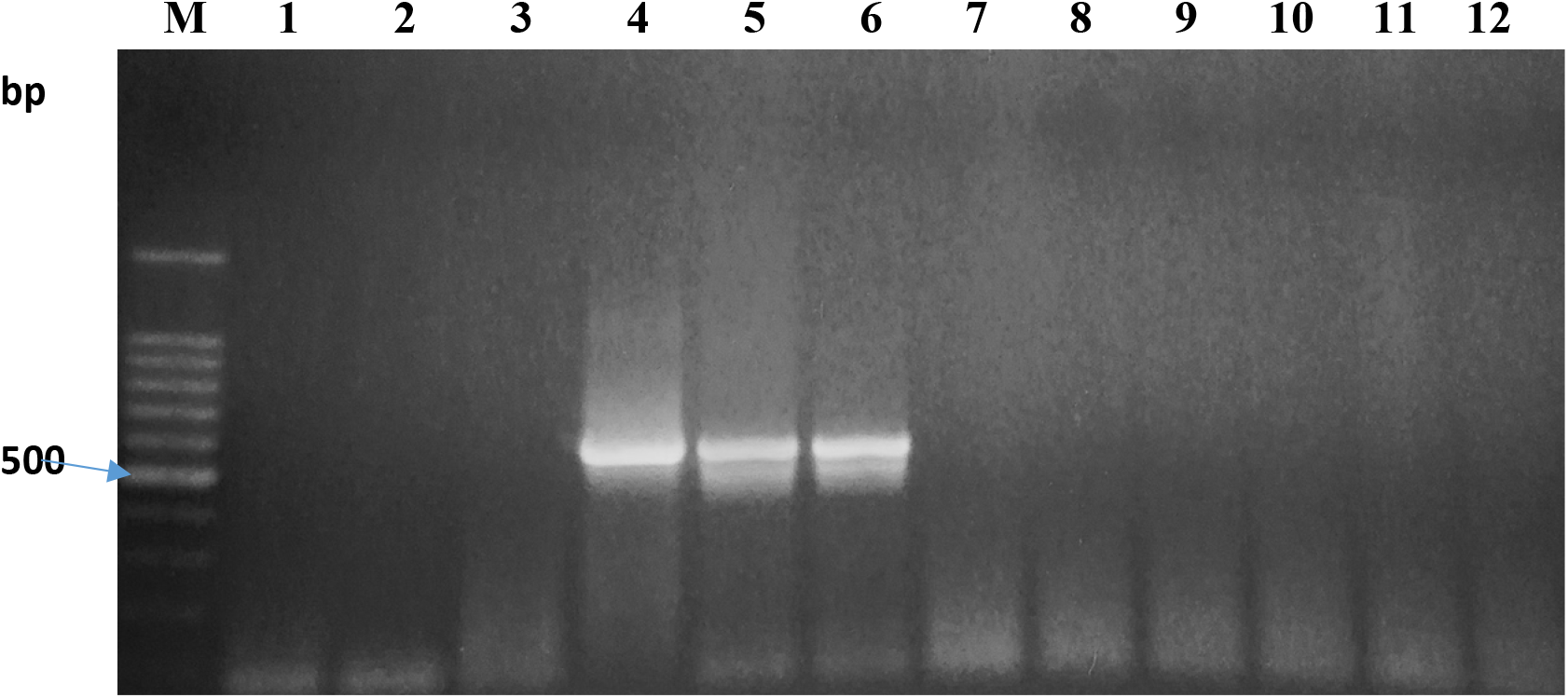
1% Agarose gel of the PCR products using *viaB* target primers and DNA templates extracted from various bacterial isolates. (Expected product size is 540 bp). M (100 bp DNA ladder). Lane 1-12 (PCR product of various bacteria using *viaB* target primers). 1-*E.* coli, 2 - *S.* boydii, 3 - *S.* Typhimurium, 4 - *S*. Typhi (s), 5 - *S*. Typhi (i), 6 - *S*. Typhi (r), 7 - *S*. Typhimurium (s), 8 - *S*. Typhimurium (i), 9 - *S*. Typhimurium (r), 10 - *S*. Enteritidis (s), 11 - *S*. Enteritidis (i), 12 - *S*. Enteritidis (r)

The *sopE* primers (SopE-Forward / SopE-Reverse) was found to be specific for all *Salmonella* spp. with PCR amplification resulting in a 465 bp product (figure 5). No amplification was observed with the *E*. coli, *S*. boydii bacterial strains.

**Figure 5.**
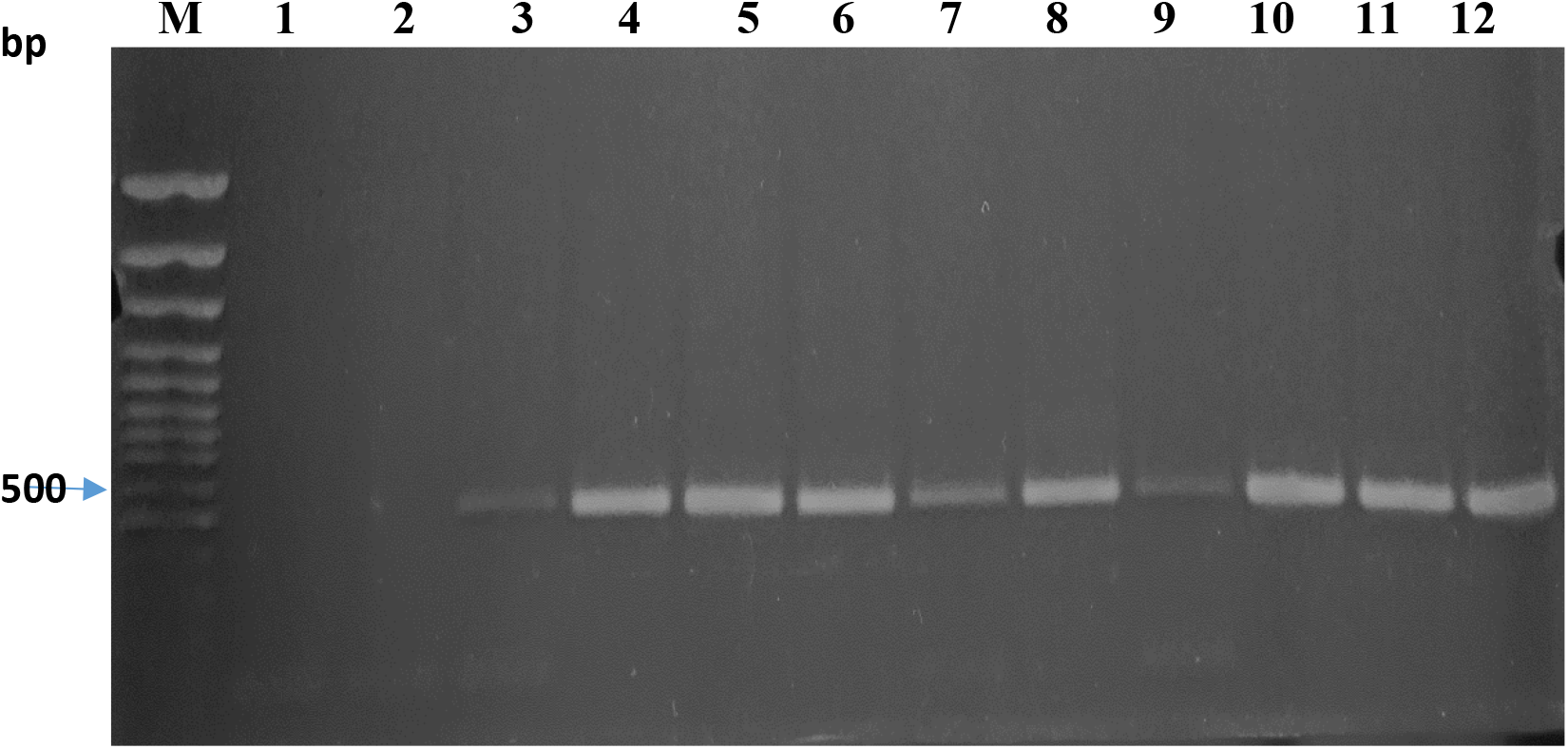
1% Agarose gel of the PCR products using sopE target primers and DNA templates extracted from various bacterial isolates. (Expected product size is 402 bp). M (100 bp DNA ladder). Lane 1-12 (PCR product of various bacteria using *sopE* target primers). 1-*E.* coli, 2 - *S.* boydii, 3 - *S.* Typhimurium, 4 - *S*. Typhi (s), 5 - *S*. Typhi (i), 6 - *S*. Typhi (r), 7 - *S*. Typhimurium (s), 8 - *S*. Typhimurium (i), 9 - *S*. Typhimurium (r), 10 - *S*. Enteritidis (s), 11 - *S*. Enteritidis (i), 12 - *S*. Enteritidis (r)

## Discussion

The goal of this study was to analyze the ability of three different sets of primers (*staA, viaB* and *sopE)* to identify *S.* Typhi strains circulating in Kenya and examine their capacity to identify and differentiate commonly isolated *Salmonella* spp. and closely related bacteria. The results demonstrated the ability of the designed primers targeting the StaA fimbrial protein gene (*staA* gene), the ViaB region DNA for Vi antigen gene (*viaB* gene) and the SopE invasion-associated secreted protein gene (*sopE* gene) to detect Salmonella serovars isolated from different regions in Kenya using conventional (qualitative) PCR. The primers targeting the *staA* gene and the *viaB* gene were both found to be specific for only *S*. Typhi while the primers targeting the SopE gene were able to detect all tested Salmonella serovars but not any of the non-Salmonella bacteria.

It has been noted that there has been an increase in multidrug-resistant (MDR) *S.* Typhi isolated from patients in Kenya since 1997 and sporadic outbreaks have also been reported in resource-poor settings, especially in slum areas [11–14]. This very alarming trend will continue to burden the Kenyan health system especially with poor diagnostic capabilities, the increasing rate of over-the-counter sale without prescription of first-line antibiotics for typhoid fever and the continued overcrowding and population rise within the slum areas [15]. The primers used in this study targeting the *staA* gene and the *viaB Salmonella* genes demonstrated their ability to only detect *S.* Typhi bacterial using conventional PCR. This may provide another tool in the accurate diagnosis of patients with typhoid fever in resource poor, endemic regions. Additionally, primers targeting the *SopE* salmonella gene may be used to also identify human nontyphoidal *Salmonella* (NTS) infections in Kenya, that have increased markedly over the years with the two main serovars isolated from cases of bacteremia and gastroenteritis with high fatality, being *S.* Typhimurium and *S.* Enteritidis [13]. With an increased prevalence of multidrug resistance among NTS serotypes [16], conventional PCR using primers targeting the *sopE* salmonella gene used in this study can be applied to identify NTS when combined together with conventional PCR using *staA* or *viaB* primers used in this study.

The *staA* primers (StaA-Forward / StaA-Reverse), and *viaB* primers (vi-Forward / vi-Reverse) used in this study have been shown to be specific only for the different strains of *S*. Typhi in Kenya and may be used as a diagnostic tool to detect the presence of the organism in individuals or foods samples using conventional pcr methods. Additionally, the *sopE* primers (SopE-Forward / SopE-Reverse) in this study were proven specific for the three most common *Salmonella* spp. while discriminating the closely related bacteria *E*. coli and *S*. boydii bacterial strains. Further studies are to be carried out with actual food samples and human samples (blood, stool or saliva) to determine the effectiveness of conventional pcr using these *staA* primers, *viaB* primers and *sopE* primers in the detection of common *Salmonella* spp. in Kenya.

## Materials and methods

### Bacterial strains

Salmonella strains were obtained from the Centre for Microbiology Research (CMR) at the Kenya Medical Research Institute (KEMRI) [17,18] and Kenyatta national hospital [14] in Nairobi, Kenya. The isolates were confirmed using slide agglutination techniques, conventional Polymerase Chain Reaction and Antimicrobial susceptibility testing. The 3 most common salmonella serovars: *Salmonella* Typhi, *Salmonella* Typhimurium and *Salmonella* Enteritidis were used in this study with each serovar consisting of 3 subtypes: susceptible (S), intermediate (I) and resistant (R). Three standard organisms: *Escherichia* coli (*E*. coli) ATCC 25922, *Shigella* boydii (*S*. boydii) ATCC 9207 and *Salmonella* Typhimurium (*S*. Typhimurium) ATCC 14022 from the National Microbiology Reference laboratory (NMRL) at the National Public Health Laboratory (NPHL) in Nairobi, Kenya were also included in the study (Table 1).

The bacteria stock cultures were maintained on a 5% nutrient broth agar slope at 4^0^C. Bacterial cultures for genomic DNA extraction were cultured in Tryptic Soy Broth (TSB) for salmonella bacteria and Luria-Bertani (LB) medium for non-salmonella bacteria. A sample of each culture was also plated on MacConkey plates for salmonella bacteria and LB agar plates for non-salmonella bacteria.

### DNA Isolation

Genomic DNA was extracted from a pure culture following the standard phenol-chloroform method [19]. Quantification of DNA was done spectrophotometrically using a UV mini 1240 UV-VIS spectrophotometer (Shimadzu, Kyoto, Japan).

### PCR Primers

1 6S rRNA target primers [20] were used for salmonella enterica bacteria confirmation. The *staA* [21], *viaB* and *sopE* primers were used to differentiate between different salmonella serovars and also differentiate non-salmonella bacteria. The list of all the primers (Inqaba biotec, Pretoria, South Africa) is indicated in table 2.

### Conventional PCR Amplification of *staA*, *viaB* and *sopE* Gene

PCR amplification of the 16S rRNA, *staA, viaB* and *sopE* genes was carried out with 50 ng of purified genomic DNA (template DNA), 1 μM of upstream primer, 1 μM of downstream primer, 1X GoTaq Green Master Mix solution (Promega, Wisconsin, USA) and nuclease-free water (Promega, Wisconsin, USA) to a final volume of 20 μl. The amplification reaction was performed on a Tprofessional thermal cycler (Biometra, Goettingen, Germany) with the following temperature and duration profile (Table 3): A volume of 10 μl of the PCR product was analyzed on a horizontal 1 % (w/v) agarose gel.

**Table 3:** Temperature programme used in the PCR amplification of the 16S rRNA, *staA*, *viaB* and *sopE* genes

| Cycle step | Temperature | Time | Number of Cycles |
| --- | --- | --- | --- |
| **Initial Denaturation** | 95 | 2 minutes | 1 |
| **Denaturation** | 95 | 1 minute |  |
| **Annealing** | 43 °C (16S rRNA)<br>51 °C (*staA*)<br>53 °C (*viaB*)<br>51 °C (*sopE*) | 45 seconds | 35 |
| **Extension** | 72 °C | 1 minute |  |
| **Final Extension** | 72 °C | 10 minutes | 1 |
| **Hold** | 4 °C | ∞ | 1 |

## Acknowledgments

We would also like to thank the Centre for Microbiology Research at the Kenya Medical Research Institute and Kenyatta National Hospital in Nairobi, Kenya for providing access to different bacterial strains and raw samples.

## Funding

This study was funded by the National Council for Science and Technology (NACOSTI) Kenya.

## Competing interests

The authors declare that they have no competing interests.

